# Analysis of subcellular transcriptomes by RNA proximity labeling with Halo-seq

**DOI:** 10.1101/2021.06.08.447604

**Authors:** Krysta L. Engel, Hei-Yong G. Lo, Raeann Goering, Ying Li, Robert C. Spitale, J. Matthew Taliaferro

**Affiliations:** Department of Biochemistry and Molecular Genetics, University of Colorado Anschutz Medical Campus; RNA Bioscience Initiative, University of Colorado Anschutz Medical Campus; Department of Chemistry, Hong Kong University; Department of Pharmaceutical Sciences, Univeristy of California, Irvine

## Abstract

Thousands of RNA species display nonuniform distribution within cells. However, quantification of the spatial patterns adopted by individual RNAs remains difficult, in part by a lack of quantitative tools for subcellular transcriptome analysis. In this study, we describe an RNA proximity labeling method that facilitates the quantification of subcellular RNA populations with high spatial specificity. This method, termed Halo-seq, pairs a light-activatable, radical generating small molecule with highly efficient Click chemistry to efficiently label and purify spatially defined RNA samples. We compared Halo-seq with previously reported similar methods and found that Halo-seq displayed a higher efficiency of RNA labeling, indicating that it is well suited to the investigation of small, precisely localized RNA populations. We then used Halo-seq to quantify nuclear, nucleolar, and cytoplasmic transcriptomes, characterize their dynamic nature following perturbation, and identify RNA sequence features associated with their composition. Specifically, we found that RNAs containing AU-rich elements are relatively enriched in the nucleus. This enrichment becomes stronger upon treatment with the nuclear export inhibitor leptomycin B, both expanding the role of HuR in RNA export and generating a comprehensive set of transcripts whose export from the nucleus depends on HuR.

## INTRODUCTION

In species ranging from yeast to mammals, many RNA species are distributed asymmetrically within cells (Cajigas et al., 2012; Lécuyer et al., 2007; Long et al., 1997; Moor et al., 2017). Mislocalization of these RNAs often results in cellular or organismal phenotypes, underscoring the importance of the process (Ephrussi and Lehmann, 1992; Ghosh et al., 2012; Long et al., 1997). RNAs are often trafficked to their destination through the action of RNA binding proteins (RBPs) that bind specific cis sequence elements, usually in the 3′ UTR, and mediate transport (Engel et al., 2020; Martin and Ephrussi, 2009). However, for most localized RNAs the identity of the cis-elements and trans-acting RBPs are unknown.

Historically, much of the work done to study RNA localization has used imaging-based approaches (Taliaferro, 2019). With these experiments, the location of RNA molecules within cells can be directly visualized. However, with the exception of new multiplexed techniques (Chen et al., 2015; Shah et al., 2016), these imaging experiments are generally limited to the interrogation of one or a few transcript species at a time. Further, short RNAs including snRNAs and snoRNAs are generally not compatible with the hybridization-based techniques used in many RNA visualization experiments.

More recently, techniques have been developed that isolate and characterize subcellular transcriptomes using high-throughput sequencing. In particular, transcriptomes of neuronal cell bodies and processes have been extensively probed in this way. These studies have relied on the elaborate, branched morphologies of neurons that facilitate their physical, mechanical separation into subcellular fractions. and have identified of hundreds of RNAs that are enriched in processes (Gumy et al., 2011; Taliaferro et al., 2016; Zappulo et al., 2017; Zivraj et al., 2010).

Asymmetric RNA distributions are also prevalent throughout cell types that lack morphologies with long, thin processes typical of neurons. Until recently, technical limitations have precluded the transcriptome-wide study of RNA trafficking in these cell types. New techniques have been developed that rely on RNA proximity to protein markers of specific subcellular locations (Fazal et al., 2019; Kaewsapsak et al., 2017; Padrón et al., 2019; Wang et al., 2019). These techniques initially label localized (i.e. protein-marker-proximal) RNA in living cells. Labeling is achieved through the enzymatic production of reactive oxygen species which diffuse from their point of generation (i.e. the localized protein marker) and leave marks on nearby RNA molecules. These marks then facilitate purification of the labeled, localized RNA away from bulk total RNA. Localized RNAs are then identified as those that are more abundant in the labeled RNA than the total RNA.

Although these proximity labeling techniques have been used to probe the RNA contents of various subcellular locations (Fazal et al., 2019; Padrón et al., 2019; Wang et al., 2019), their enzymatic approaches to radical generation may limit the amount of radicals produced and therefore the sensitivity of RNA labeling. A nonenzymatic technique for radical generation using a light-sensitive Halo ligand fluorophore, dibromofluorescein (DBF) was recently described (Li et al., 2017, 2018), but its ability to provide comprehensive characterization of subcellular transcriptomes has not been tested. Here, we demonstrate that DBF-mediated RNA proximity labeling can be used to efficiently characterize subcellular transcriptomes and derive mechanistic insights into factors controlling their composition.

## RESULTS

### Halo-seq allows in situ alkynylation of spatially resolved RNA populations

Halo-seq uses HaloTag domains genetically fused to a protein that specifically marks the subcellular location of interest. HaloTags are protein domains that covalently bind to a class of small molecules called Halo ligands (Los et al., 2008). If a Halo ligand is added to cells expressing a spatially restricted protein containing a HaloTag, the ligand will therefore be similarly restricted.

The Halo ligand used in Halo-seq is dibromofluorescein (DBF). DBF generates singlet oxygen radicals when irradiated with green light (Li et al., 2017, 2018). The high reactivity of these radicals restricts their diffusion from the DBF source to a radius of approximately 100 nm (**Figure 1A**) (Li et al., 2017; Moan, 1990; Skovsen et al., 2005). When these radicals react with an RNA base within this 100 nm radius, the base becomes oxidized and prone to nucleophilic attack by a cell-permeable, alkyne-containing nucleophile, propargylamine (PA). RNA molecules within 100 nm of a DBF molecule are therefore selectively alkynylated. Importantly, this alkynylation occurs while the cell is alive and intact, providing confidence that the alkynylated RNA molecules are spatially coincident with the region of interest.

**Figure 1.**
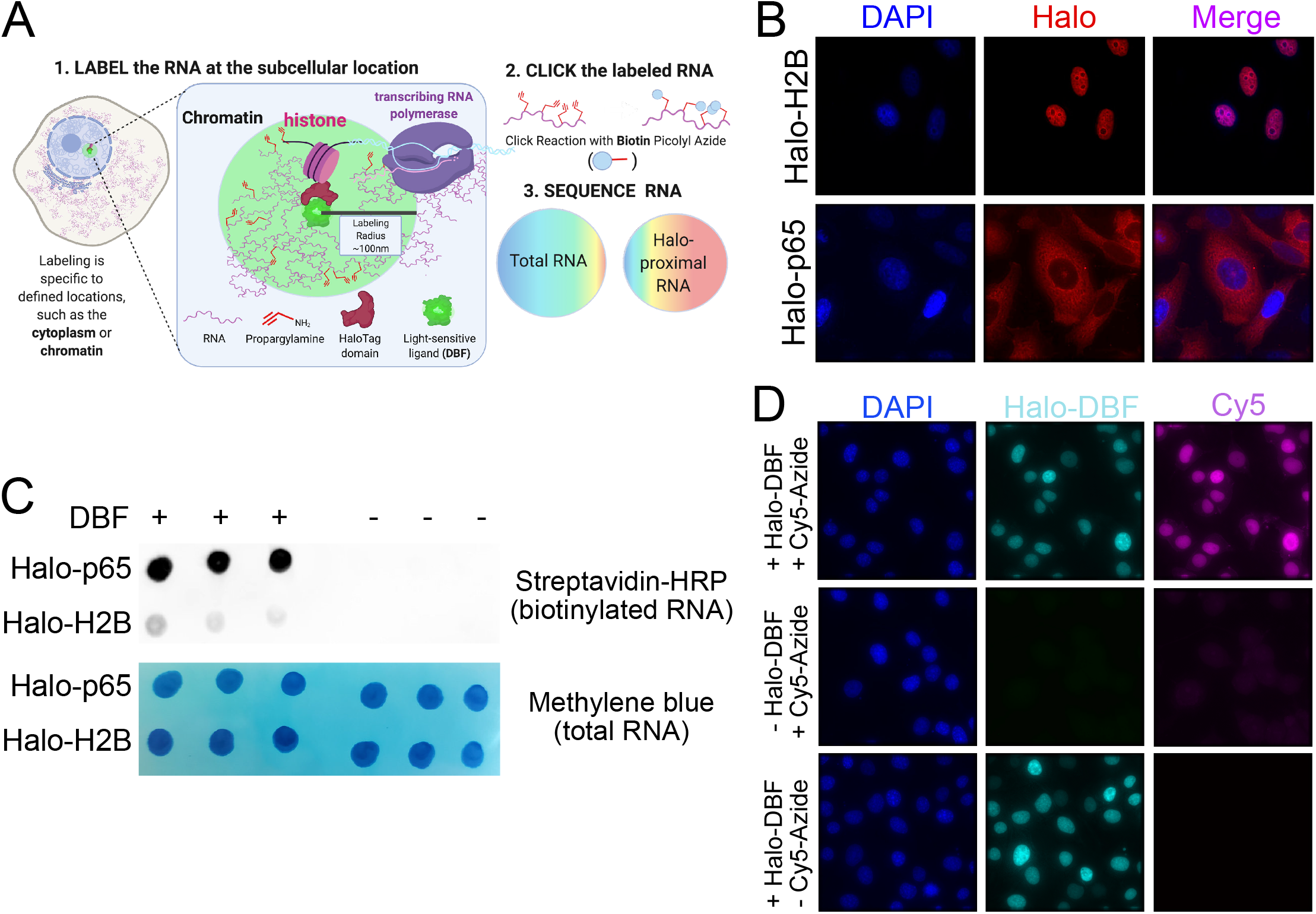
Halo-seq facilitates the biotinylation of RNA transcripts in proximity to spatially restricted Halo-DBF molecules. (A) Overview of the Halo-seq procedure. A HaloTag protein domain is genetically fused to a protein which localizes to a subcellular location of interest. Because Halo ligands specifically bind HaloTags, this also spatially restricts a DBF Halo ligand. When irradiated with green light, DBF emits oxygen radicals that label nearby RNAs, resulting in their alkynylation. Alkynykated RNAs are substrates for *in vitro* biotinylation using “Click” chemistry, which allows the localized RNAs to be separated from the bulk RNA sample with streptavidin pulldown, and quantified using high-throughput sequencing. (B) HaloTag protein domains fused to histone H2B and p65 are localized to chromatin, and the cytoplasm, respectively. HaloTag domains are visualized through the addition of a Halo ligand fluorophore. (C) RNA samples taken from cells expressing Halo fusions can be biotinylated *in vitro*, and this biotinylation is dependent upon the addition of a DBF Halo ligand to cells. (D) Alkynylated molecules can be visualized *in situ* by fusing them with fluorophores (e.g. Cy5-azide) using Click chemistry. Alkynylated molecules are restricted to the nucleus in cells containing H2B-Halo. They are only detectable in cells treated with both DBF and Cy5-azide, demonstrating the ability of HaloTag-restricted DBF to induce alkynylation of biomolecules.

Following the in-cell alkynylation, cells are lysed and total RNA is collected. Alkynylated molecules are efficient substrates for *in vitro* Cu(I)-catalyzed azide-alkyne cycloaddition (CuACC) “Click” chemistry in which alkyne-containing molecules and azide-containing molecules are linked (Hein et al., 2008). Reaction with biotin-azide therefore selectively biotinylates RNA molecules that were in close proximity to DBF, facilitating their isolation with streptavidin and analysis by high-throughput sequencing. By comparing RNA abundances in total RNA samples and streptavidin-purified samples, transcripts that were enriched in the subcellular region of interest are identified.

To test the ability of Halo-seq to purify and quantify subcellular transcriptomes, we restricted HaloTag domains to chromatin and the cytoplasm. These HaloTag localizations were achieved by fusing to histone H2B and p65, a cytoplasmically-localized NF-kappa B subunit, respectively. Restriction of Halo and DBF to these locations was previously demonstrated to enable RNA tagging with high spatial resolution as assayed by RT-qPCR (Li et al., 2017, 2018). We reasoned that since the RNA contents of nucleic and cytoplasmic compartments have been well-characterized, we would be able to easily assay the accuracy of the Halo-seq procedure. We integrated a single, doxycycline-inducible copy of these fusion constructs into the genomes of HeLa cells using cre-mediated recombination. We then verified the proper subcellular localization of the fusion proteins by visualizing them with fluorescent Halo ligands (**Figure 1B**).

We next assayed the ability of Halo-seq to produce biotinylated RNA using an RNA dot blot in which biotinylated RNA is detected using streptavidin-HRP. We found that RNA samples from both the Halo-p65 and H2B-Halo expressing lines could be efficiently biotinylated (**Figure 1C**). This biotinylation was dependent upon the addition of DBF to the cells, indicating that RNA biotinylation is dependent on the Halo-seq procedure and not due to an endogenous activity. Since all samples were subjected to the same procedures *in vivo* and *in vitro*, this also indicates that the level of biotinylation via any background labeling *in vitro* is very low.

Next, we sought to visualize the localization of alkynylated molecules in cells following treatment with DBF and excitation with green light. This can be accomplished by fixing the cells after alkynylation and performing the Click reaction in situ with an azide coupled to a fluorescent molecule. With this approach, we found that the nucleus of cells expressing H2B-Halo was rich in alkynes, as evidenced by strong nuclear Cy5-azide signal that was dependent on the addition of DBF (**Figure 1D**). Overall, these results demonstrate that our approach herein is high-resolution and enables imaging based analysis of tagging.

### Quantification of subcellular transcriptomes with Halo-seq

We then characterized nuclear and cytoplasmic transcriptomes by comparing RNA abundances in samples taken before and after streptavidin-pulldowns. RNAs that were enriched at the subcellular location of interest should be enriched in the streptavidin-pulldown sample relative to the input sample. For each condition, either three or four biological replicates were used. We prepared rRNA-depleted libraries for high-throughput sequencing, quantified transcript expression in the resulting data using Salmon (Patro et al., 2017), and collapsed the data to gene-level abundances with tximport (Soneson et al., 2015). Genes whose abundances were significantly different between input and streptavidin-pulldown samples were identified using DESeq2 (Love et al., 2014).

Principal component analysis of gene expression profiles revealed that input and pulldown samples were markedly different from each other in both the H2B and p65 experiments (**Figure S1A, B**), suggesting that the Halo-seq procedure reproducibly isolated distinct RNA populations. Hundreds of genes were both enriched and depleted in the pulldowns from the H2B and p65 experiments (**Figure 2A,B, Tables S1 and S2**), further indicating that Halo-seq can identify distinct nuclear and cytoplasmic transcriptomes.

**Figure 2.**
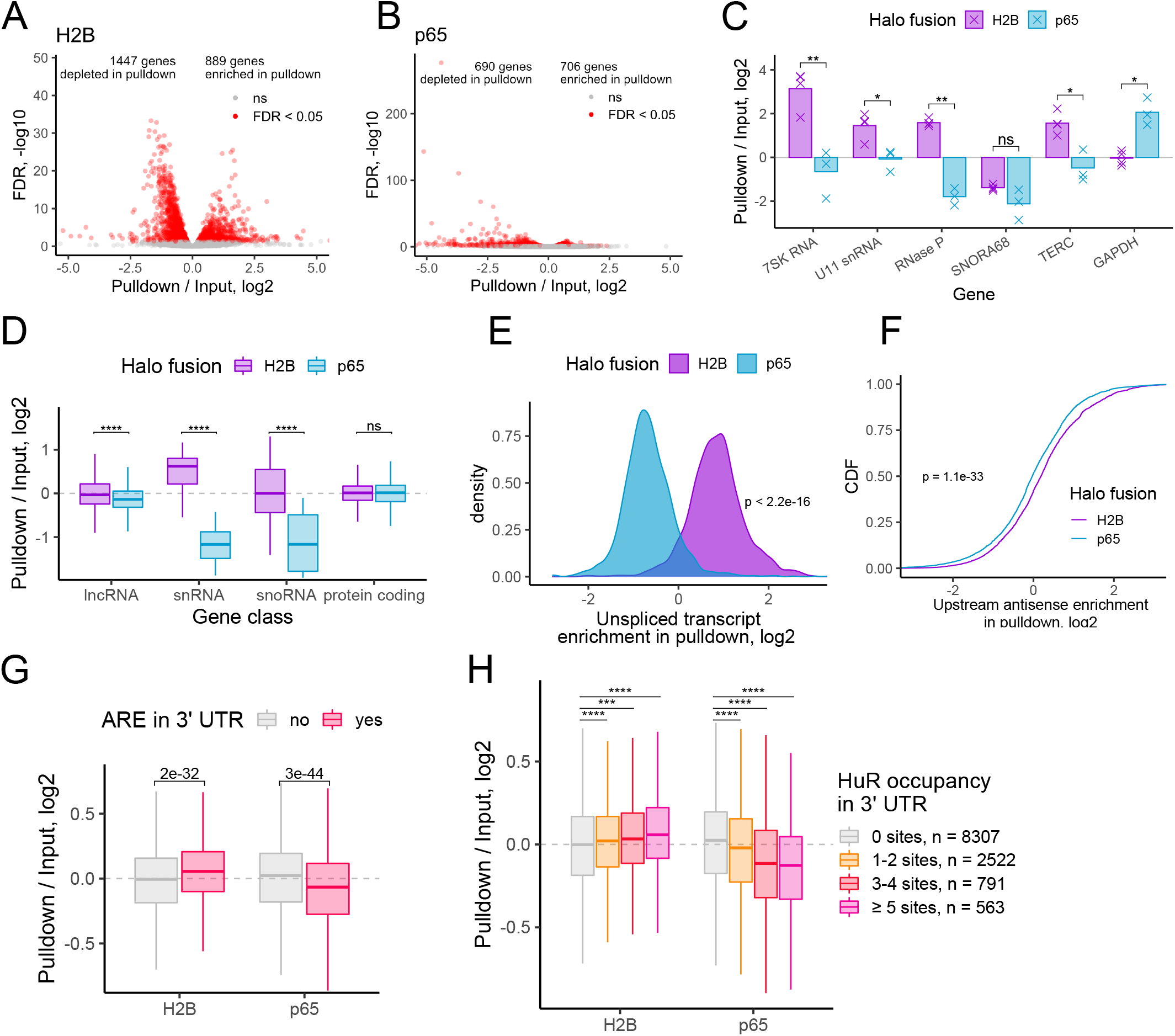
Halo-seq quantification of nuclear and cytoplasmic transcriptomes. (A) Differentially expressed genes in a comparison of pulldown and input RNA samples following Halo-seq RNA labeling using a H2B-Halo fusion. (B) As in A, but using a Halo-p65 fusion. (C) Halo-seq enrichments for selected RNA species known to be localized to the nucleus or cytoplasm. (D) Halo-seq enrichments for defined classes of RNAs. (E) Halo-seq enrichments for unspliced, intron-containing transcripts. The ratio of the abundance of unspliced transcripts to spliced transcripts was calculated for each gene. These ratios were then compared in input and pulldown samples. (F) Halo-seq enrichments for promoter-proximal upstream antisense transcripts. The ratio of the abundance of promoter-proximal upstream antisense to downstream transcripts was calculated for each gene. This ratio was then compared in input and pulldown samples. (G) Genes were binned by whether or not their 3′ UTR contained an AU-rich element (ARE). Gene enrichments in Halo-seq pulldown and input samples were then compared. (H) Genes were binned by the number of HuR binding sites (as defined by CLIP-seq) in their 3′ UTRs. Gene enrichments in Halo-seq pulldown and input samples were then compared. All significance tests were performed using a Wilcoxon rank-sum test. p value notation: * < 0.05, ** < 0.01, *** < 0.001, **** < 0.0001.

We then further analyzed the H2B and p65 samples, paying special attention to RNA species and classes known to be enriched in either the nuclear or cytoplasmic compartments. Several RNA species, including the 7SK RNA, RNase P, and TERC, function primarily in the nucleus and are known to accumulate there (Altman, 2011; Egloff et al., 2018; Zhang et al., 2011). Accordingly, all of these RNAs were enriched in the pulldown of the H2B experiment and depleted in the pulldown of the p65 experiment (**Figure 2C**). Conversely, GAPDH mRNA is known to accumulate to a higher relative level in the cytoplasm than in the nucleus, and this was also reflected in the relative enrichments from the H2B and p65 experiments (**Figure 2C**). Small nucleolar RNAs (snoRNAs), a class of small noncoding RNAs involved in ribosomal RNA maturation and ribosome biogenesis, are primarily localized to the nucleolus (Liang et al., 2019). Accordingly, the snoRNA SNORA68 was depleted in both the H2B and p65 pulldowns. Because the nucleus and nucleolus are in close proximity to each other yet H2B is excluded from the nucleolus, this indicates that RNA labeling with Halo-seq displays a high degree of spatial selectivity.

To generalize these results, we then compared enrichment for classes of RNAs. Long noncoding RNAs (lncRNAs) are generally depleted from the cytoplasm (Lubelsky and Ulitsky, 2018; Shukla et al., 2018). In agreement with this, we observed that lncRNAs, as a class, were significantly less enriched in the p65 pulldown than in the H2B pulldown (**Figure 2D**). Similarly, snRNAs are enriched in the nucleus (Matera and Wang, 2014), and Halo-seq found them to be enriched in the H2B pulldown and depleted in the p65 pulldown. This contrasts with protein-coding RNAs, which as a class spend substantial time in both the nuclear and cytoplasmic compartments. These RNAs were equally enriched in the H2B and p65 pulldowns.

We then turned to the quantification of dynamically processed RNA species. Most protein-coding human RNAs undergo splicing in the nucleus before export to the cytoplasm, and for many RNAs this splicing happens cotranscriptionally (Herzel et al., 2017). We therefore wondered if we could observe an enrichment for unspliced pre-mRNA in close proximity to chromatin via the H2B pulldown. For each gene, we quantified the relative abundance of spliced and unspliced RNA in each sample using a custom Salmon index that contained both spliced and unspliced species. We then compared the ratio of unspliced to spliced abundances in the input and pulldown samples. For the H2B samples, we found that unspliced transcripts were relatively enriched in the pulldown compared to the input, whereas for the p65 samples, unspliced transcripts were depleted in the pulldown (**Figure 2E, Figure S1C,D**), demonstrating that Halo-seq is able to interrogate the localization of dynamically processed RNAs.

Next, we asked whether Halo-seq could quantify intrinsically unstable and short-lived RNAs. During transcription initiation, antisense RNAs are produced from the region upstream of the promoter (Flynn et al., 2011). These upstream antisense transcripts are highly unstable and very unlikely to leave the nucleus. We reasoned therefore that they should be enriched in the H2B pulldown relative to the p65 pulldown. For each gene, we calculated the ratio of upstream antisense RNAs to downstream sense RNAs. We then compared this ratio in input and pulldown samples. We found that upstream antisense RNAs were significantly more enriched in the H2B pulldown than the p65 pulldown (**Figure 2F**), indicating that Halo-seq can quantify highly unstable RNAs with spatial precision.

### HuR-bound transcripts are enriched in the nucleus and depleted in the cytoplasm

Given the assurance in the quality of our spatially resolved nuclear and cytoplasmic transcriptomes, we searched for RNA features that discriminated between nucleus- and cytoplasm-enriched transcripts. Interestingly, we found that genes that contained AU-rich elements (AREs) in their 3′ UTRs were enriched in the H2B pulldown and depleted from the p65 pulldown (**Figure 2G**). AREs are RNA sequence motifs, often located in 3′ UTRs, that are most often thought to regulate transcript stability, but have also been noted to regulate RNA export, but the mechanism and the RNA binding selection are unclear (Chen and Shyu, 1995; Gallouzi and Steitz, 2001). HuR is an RNA-binding protein known to bind AREs (Lebedeva et al., 2011). Accordingly, we found a dose-dependent relationship between the number of CLIP-defined HuR binding sites (Mukherjee et al., 2011) in a gene’s 3′ UTR and the degree of its enrichment in the H2B pulldown. Conversely, we found an inverted dose-dependent relationship between 3’ UTR HuR binding sites and its depletion in the p65 pulldown (**Figure 2H**). These results suggest that specific RNA sequences and RBPs can modulate the relative nuclear and cytoplasmic abundances of a transcript (Yi et al., 2010).

### Halo-seq distinguishes transcriptomes of compartments in close proximity to each other

Although the results from our analysis of nuclear and cytoplasmic transcriptomes were highly encouraging, these two cell compartments are separated by a well-defined membrane and therefore may not provide the best scenario with which to test the spatial specificity of Halo-seq. We reasoned instead that a more rigorous test of the technique would be to differentiate the RNA contents of non-membrane-bound organelles. We therefore chose to interrogate another subcellular compartment, the nucleolus.

We began by fusing a HaloTag to a specific marker of nucleoli, Fibrillarin (Ochs et al., 1985). This fusion specifically localized to nucleoli and was distinguishable from the chromatin-associated H2B-Halo fusion using fluorescence imaging (**Figure 3A**). We performed the Halo-seq alkynylation and biotinylation procedures and detected biotinylated RNA from Halo-Fibrillarin-expressing cells that was dependent upon the addition of DBF, indicating that nucleolar RNA was being labeled by Halo-seq (**Figure 3B)**. We then sequenced the transcripts enriched by Halo-Fibrillarin using libraries created with rRNA depletion. Gene expression values of streptavidin input and pulldown samples were well separated by PCA, suggesting that the Fibrillarin-mediated labeling occurred on specific transcripts (**Figure S2A**). Halo-Fibrillarin labeled RNA was enriched for 890 genes and depleted for 823 genes (FDR < 0.05), indicating that the nucleolus contains an RNA population distinct from bulk cellular RNA (**Figure 3C, Table S3**).

**Figure 3.**
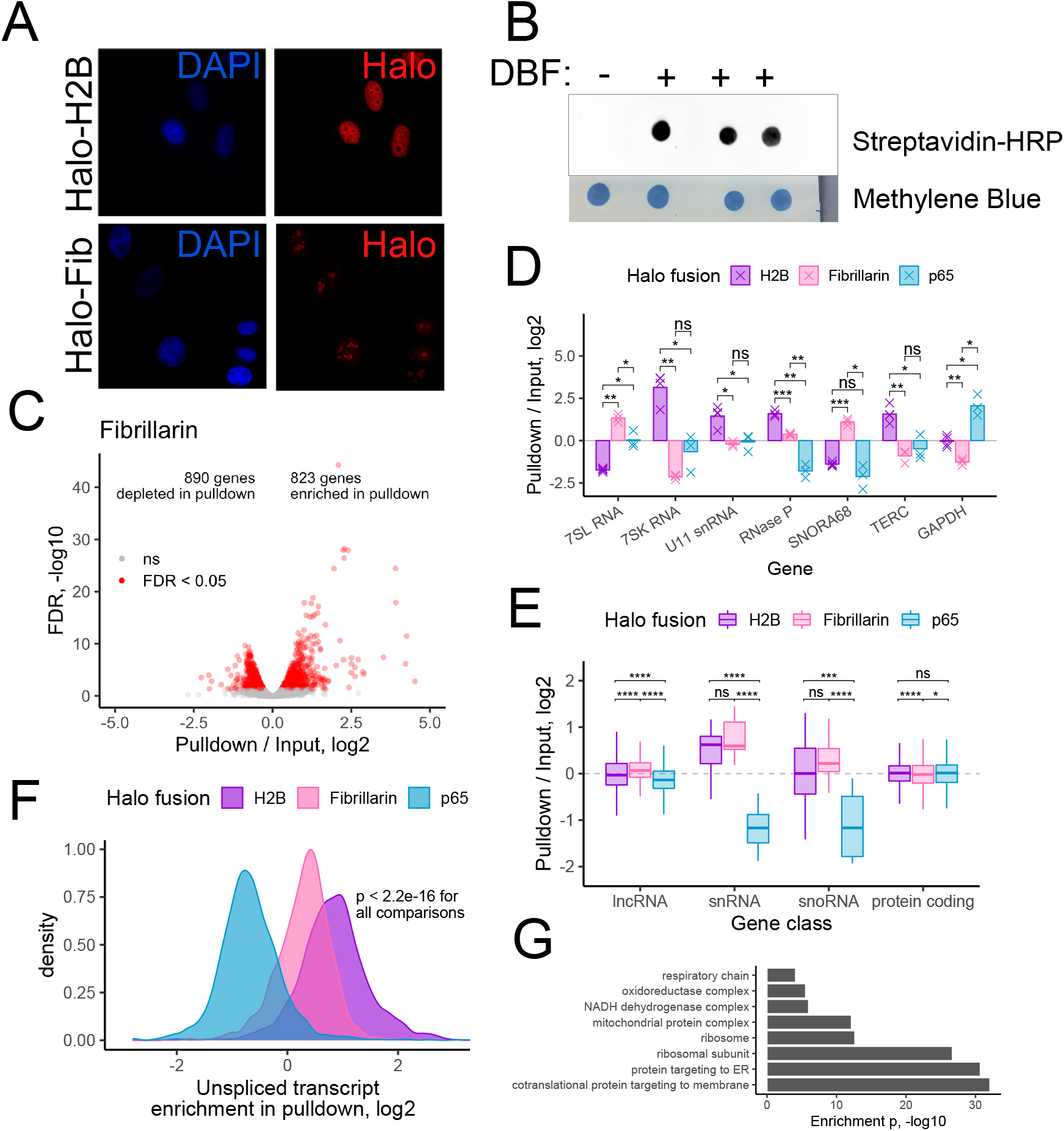
Halo-seq quantification of the nucleolar transcriptome. (A) Visualization of the subcellular location of histone H2B and fibrillarin Halo domain fusions. Halo fusion proteins were visualized using a fluorescent Halo ligand. (B) RNA dot blot of RNA collected from cells expressing a Halo-fibrillarin fusion protein. (C) Differentially expressed genes in a comparison of pulldown and input RNA samples following Halo-seq RNA labeling using a Halo-fibrillarin fusion. (D) Halo-seq enrichments for selected RNA species known to be localized to the nucleus, nucleolus, or cytoplasm. (E) Halo-seq enrichments for defined classes of RNAs. (F) Halo-seq enrichments for unspliced, intron-containing transcripts. As in Figure 2E, the ratio of the abundance of unspliced transcripts to spliced transcripts was calculated for each gene. This ratio was then compared in input and pulldown samples. (G) Enriched gene ontology terms derived from RNAs identified as localized to the nucleolus. All significance tests were performed using a Wilcoxon rank-sum test. p value notation: * < 0.05, ** < 0.01, *** < 0.001, **** < 0.0001.

As with the H2B and p65 experiments, we looked at the enrichment of RNA species previously known to be present in nucleoli (**Figure 3D**). Encouragingly, we found that 7SL RNA was enriched in the Fibrillarin pulldown. This RNA is part of the signal recognition particle (SRP) ribonucleoprotein complex. SRP is involved in protein targeting to the endoplasmic reticulum in the cytoplasm, but it is assembled in the nucleolus (Politz et al., 2000). Similarly, we found that the snoRNA SNORA68 was enriched in the Fibrillarin pulldown but not in either the nuclear or the cytoplasmic experiments. This is an encouraging result since snoRNAs are known to be primarily localized to the nucleolus (Liang et al., 2019). Importantly, both of these RNAs were depleted from the H2B pulldown, indicating that Halo-seq can distinguish the chromatin-proximal and nucleolar transcriptomes even though these two compartments are in close proximity to each other. Conversely, the transcription-regulating 7SK RNA and the telomere-regulating TERC RNA were enriched in the H2B pulldown and depleted in the Fibrillarin pulldown, further highlighting the spatial specificity of Halo-seq.

To generalize these results, we calculated the enrichments of classes of RNAs (**Figure 3E**). snRNAs were depleted from the cytoplasmic p65 pulldown but were similarly enriched in the H2B and Fibrillarin pulldowns. Although snRNAs are mainly thought of in the context of chromatin-associated pre-mRNA splicing, they do transiently pass through the nucleolus and are modified there (Gerbi and Lange, 2002; Yu et al., 2001). snoRNAs as a class were significantly more enriched in the Fibrillarin pulldown than the H2B pulldown, again highlighting the spatial specificity of Halo-seq. Unspliced pre-mRNAs were significantly less enriched in the fibrillarin pulldown than the H2B pulldown (**Figure 3F**), further highlighting the specificity.

A gene ontology analysis of RNAs enriched in the Fibrillarin pulldown revealed many terms associated with ribosome biogenesis, as would be expected with this being a well-characterized of the nucleolus (**Figure 3G, S2B**). Interestingly, we also observed an enrichment of nuclearly-encoded mitochondrial genes in the Fibrillarin pulldown. This observation is particularly intriguing as mitochondrial and nucleolar proteins have been observed to colocalize more often than expected (Thul et al., 2017).

### Comparison of Halo-seq to other high-throughput methods for studying RNA localization

Other methods for the purification and analysis of subcellular transcriptomes have been previously reported (Benoit Bouvrette et al., 2018; Fazal et al., 2019; Padrón et al., 2019; Wang et al., 2019). We therefore set out to compare the sensitivity and specificity of Halo-seq to these methods.

Halo-seq utilizes a dibromofluorescein (DBF) Halo ligand; however, this ligand is not commercially available. We therefore tested the ability of this ligand to induce RNA labeling in comparison to the more commonly used and commercially available fluorescein Halo ligand. RNA biotinylation was assayed using dotblots following reactions with these two ligands in HaloTag-expressing cells (**Figure 4A**). RNA labeling reactions with DBF produced considerably more biotinylated RNA than those with fluorescein, indicating that DBF is more efficient at facilitating RNA alkynylation than fluorescein. This is consistent with previous reports demonstrating that DBF is a more efficient singlet oxygen generator and has a much higher singlet oxygen yield (ΦΔ; 0.42) than that of fluorescein (Gandin et al., 1983). Critically, the increased singlet oxygen production by DBF results in higher yield of RNA oxidation for subsequent tagging by propargylamine. This difference is explained by heavy atoms (bromine) increasing the likelihood that photoexcited chromophores will undergo an intersystem crossing (ISC) to the triplet state, from which ^1^O_2_ can be generated from ground state O_2_. The drastic difference in RNA tagging suggests that the choice of fluorophore and understanding its singlet oxygen yield is critical for the further development of Halo-seq.

**Figure 4.**
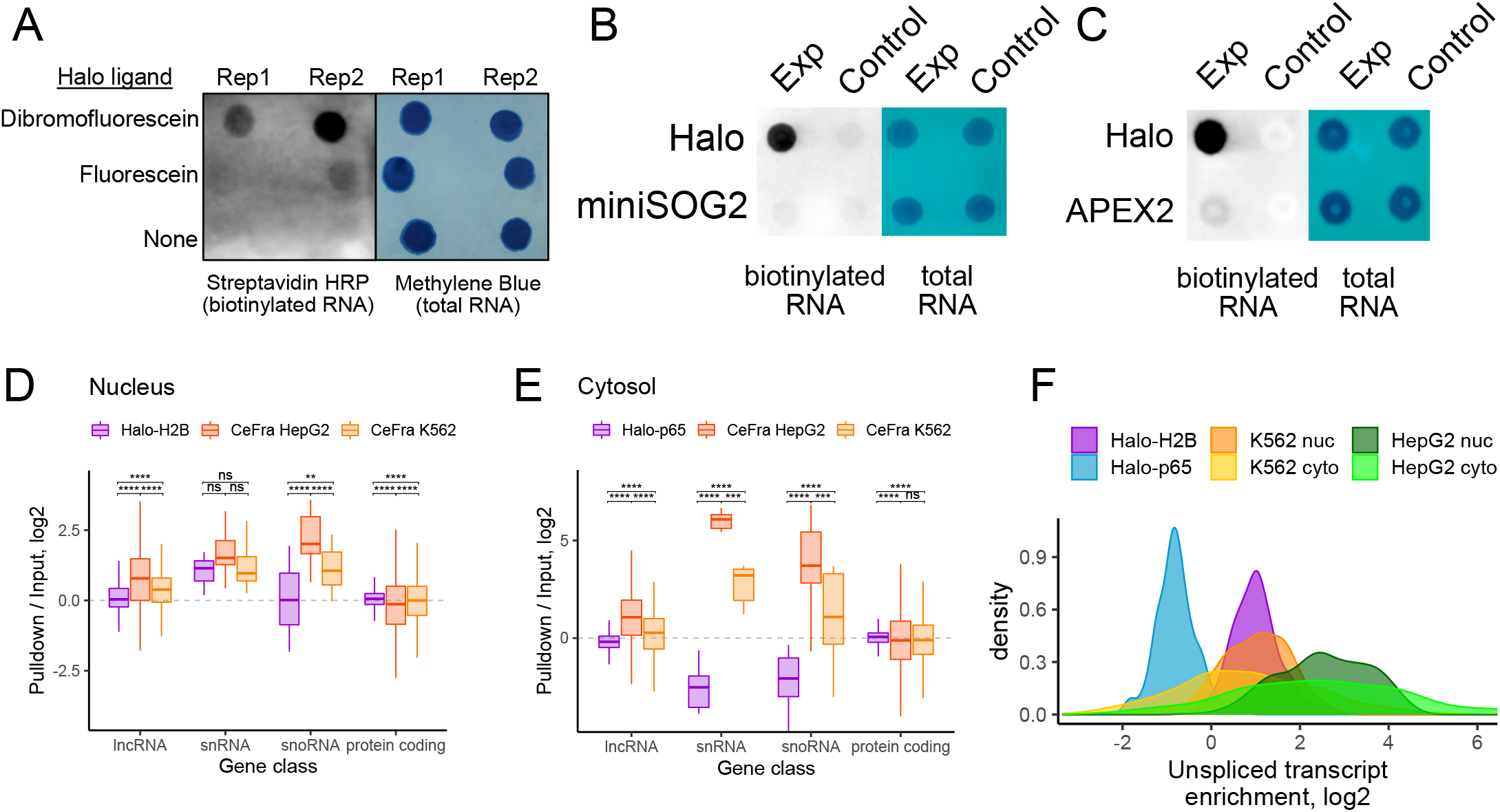
Comparison of Halo-seq with similar methods for quantifying RNA localization transcriptome-wide. (A) RNA biotinylation, as assayed by RNA dot blot, in cells expressing a HaloTag domain and treated with Halo ligand forms of dibromofluorescein (DBF) or fluorescein. (B) Comparison of RNA biotinylation efficiency of Halo-seq and CAP-seq, a similar RNA proximity labeling method that uses miniSOG2, an enzymatic singlet oxygen generator. For the Halo-expressing cells, the experimental and control conditions correspond to the addition or omission of DBF, respectively. For the miniSOG2-expressing cells, the experimental and control conditions correspond to the treatment or omission of blue light irradiation. (C) Comparison of RNA biotinylation efficiency of Halo-seq and APEX-seq. For the Halo-expressing cells, the experimental and control conditions correspond to the addition or omission of DBF, respectively. For the APEX2-expressing cells, the experimental and control conditions correspond to the inclusion or omission of hydrogen peroxide. (D) Comparison of nuclear enrichments for selected gene classes using data produced by Halo-seq and CeFra-seq. The Halo-seq data corresponds to enrichments from the H2B-Halo-mediated labeling experiment described previously. The CeFra-seq data corresponds to enrichments in a biochemically defined nuclear fraction compared to total RNA from an unfractionated sample. (E) As in D, comparison of cytosolic enrichments for selected gene classes. The Halo-seq data corresponds to enrichments from the Halo-p65-mediated labeling experiment described earlier. The CeFra-seq data corresponds to enrichments in a biochemically defined cytosolic fraction compared to total RNA from an unfractionated sample. (F) Enrichments for unspliced, intron-containing transcripts. The ratio of the abundance of unspliced transcripts to spliced transcripts was calculated for each gene. For Halo-seq samples, this ratio was then compared in input and pulldown samples. For CeFra-seq samples, this ratio was then compared in fractionated (either nuclear or cytosolic) and total, unfractionated samples. All significance tests were performed using a Wilcoxon rank-sum test. p value notation: * < 0.05, ** < 0.01, *** < 0.001, **** < 0.0001.

Cap-seq, a similar method that uses the singlet oxygen generator miniSOG2 to label RNAs was recently reported (Wang et al., 2019). In this approach, the light-activated protein miniSOG2 generates radicals that, together with propargylamine, can alkynylate nearby RNA molecules. To test the relative labeling efficiency of the miniSOG2- and Halo-based methods, we created HeLa cell lines containing either miniSOG2 or Halo constructs integrated at a single, defined locus. Importantly, both of these constructs contained HA epitope tags, allowing the direct comparison of their expression levels using immunoblotting with an anti-HA antibody. We performed the alkynylation reactions for both samples, irradiating the Halo sample with green light, and the miniSOG2 sample with a previously reported optimized blue light wavelength (Wang et al., 2019).

As expected, we observed robust RNA biotinylation in the Halo sample that was dependent upon the addition of DBF to the cells (**Figure 4B**). In contrast, we observed relatively little RNA biotinylation in the miniSOG2 samples, both with and without irradiation with blue light. In these tests, the Halo protein was more highly expressed than the miniSOG2 protein, even though their respective constructs used the same promoter and were integrated into the same genomic locus (**Figure S3A**). Nevertheless, the difference in RNA biotinylation efficiency between the methods (>100 fold) could not be explained by the differences in protein expression (approximately 4 fold). One explanation for the striking difference between the two is the singlet oxygen yield, which is related to the stability of both the chromophore and its excited state. DBF singlet oxygen yield is 0.42 and miniSOG is reported to be 0.03 (Ruiz-González et al., 2013). It is also well established in miniSOG that the formation of radicals (and singlet oxygen) results in protein damage near the chromophore, which limits the generation of more radicals over time. Prolonged irradiation to blue light leads to several structural alterations of miniSOG (Torra et al., 2019), which include photodegradation of FMN oxidation of the quenching side chains. These results are expected to be a general feature of flavin-binding proteins (Endres et al., 2018).

Another RNA proximity labeling technique that utilizes radical generation is APEX-seq (Fazal et al., 2019; Padrón et al., 2019). Similarly to Halo-seq, APEX-seq relies on a spatially restricted fusion protein to facilitate spatially restricted RNA labeling. In APEX-seq, a spatially restricted protein is fused to the radical-generating enzyme APEX2. We compared the efficiency of DBF-mediated and APEX2-mediated RNA labeling by creating a cell line that expressed a Halo-APEX2 fusion protein. This strategy ensures that the HaloTag domain and APEX2 enzyme are expressed at equal amounts (**Figure S3B**). Using these cells, we then performed either the Halo-seq procedure or APEX-seq (Fazal et al., 2019; Padrón et al., 2019). With both methods, we observed more labeling in the experimental samples than their respective controls (DBF omission for Halo-seq, hydrogen peroxide omission for APEX-seq). However, we observed approximately 10-fold more labeling in the Halo-seq experimental sample than the APEX-seq experimental sample, suggesting that DBF-mediated RNA labeling is more efficient than APEX2-mediated RNA labeling (**Figure 4C**). Differences in tagging may be due to the short-lived nature of the generated radicals and their known propensity to cleave RNA, which would reduce the reactive capacity of generated tyramide radicals for tagging.

To further compare Halo-seq and APEX-seq, we compared RNA enrichments (pulldown / input) in cytoplasmic, nuclear, and nucleolar samples using hierarchical clustering (**Figure S3C**). Unfortunately, APEX-seq RNAseq libraries were produced using polyA-enrichment, precluding quantification of the small RNA species (e.g. snRNAs and snoRNAs) that can serve as specific markers of these subcellular fractions. Nevertheless, we found that cytoplasmic samples from Halo-seq and APEX-seq clustered together, indicating their similarity. Nuclear samples from the two methods also clustered together. Conversely, the nucleolar samples from the two methods did not cluster together. Nucleolar samples from APEX-seq were similar to the nuclear samples from both methods while the nucleolar samples from Halo-seq were well separated from all other samples. However, it is not possible to determine whether these differences are due to differences in spatial selectivity or library preparation method.

Other methods to study RNA localization rely on biochemical fractionation rather than proximity labeling (Benoit Bouvrette et al., 2018). One such study used CeFra-seq to biochemically separate two cultured cell lines, HepG2 and K562, into nuclear, cytosolic, insoluble, and membrane-associated fractions. RNAseq libraries prepared from these fractions were constructed using rRNA-depletion, allowing direct comparison of these results with Halo-seq. We first used hierarchical clustering to compare gene enrichment values (pulldown / input for Halo-seq, fraction / total for CeFra-seq) produced by the two methods (**Figure S3D**).

While the nuclear samples of the two methods were well correlated, the cytoplasmic CeFra-seq samples were more related to the nucleolar Halo-seq samples than the cytoplasmic Halo-seq samples.

We then compared gene enrichments from specific gene classes. Whereas both methods showed enrichments for snRNAs in their nuclear fraction, CeFra-seq showed enrichments for snoRNAs in the nuclear fraction while Halo-seq did not (**Figure 4D**). This is likely due to the inability of CeFra-seq to distinguish between the nucleus and nucleolus. Interestingly, in the cytoplasmic CeFra-seq samples, snRNAs and snoRNAs were enriched, even though these are nuclear and nucleolar markers, respectively (**Figure 4E**). Similarly, unspliced transcripts were comparably enriched in the nuclear and cytoplasmic CeFra-seq samples even though unspliced transcripts are generally found in the nucleus (**Figure 4F**). From these findings, we conclude that Halo-seq is better suited to isolating subcellular transcriptomes than the biochemical CeFra-seq method. Further, the flexibility of Halo-seq allows the interrogation of subcellular compartments that are not amenable to biochemical purification.

### Halo-seq identifies RNA targets of the CRM1-dependent RNA export pathway

Given that we had established that Halo-seq allows the efficient and accurate profiling of subcellular transcriptomes, we moved to the quantification of RNA localization following perturbation. A subset of transcripts in mammalian cells depend upon the action of CRM1 for export from the nucleus, and the small molecule leptomycin B (LMB) inhibits the action of CRM1 (Okamura et al., 2015). While the identity of a handful of RNAs that are nuclearly retained following LMB treatment are known (Brennan et al., 2000; Jang et al., 2003), the full complement of RNAs that depend on CRM1 for export remains unknown.

To more fully investigate the dependence on CRM1 for RNA export, we identified nuclearly enriched RNAs using Halo-seq with a H2B-Halo fusion in the presence and absence of LMB. Following LMB treatment, we expected an increased nuclear abundance for a subset of RNAs whose nuclear export depends on CRM1.

We first verified that the nuclear localization of the H2B-Halo fusion protein did not change following LMB treatment (**Figure 5A**). Our cytoplasmic Halo-p65 marker does contain a CRM1-dependent nuclear export motif and was correspondingly restricted to the nucleus following LMB treatment (**Figure 5A**). While this excludes it from being a useful cytoplasmic marker in this experiment, it does provide clear validation that the LMB treatment inhibited CRM1 activity under these conditions.

**Figure 5.**
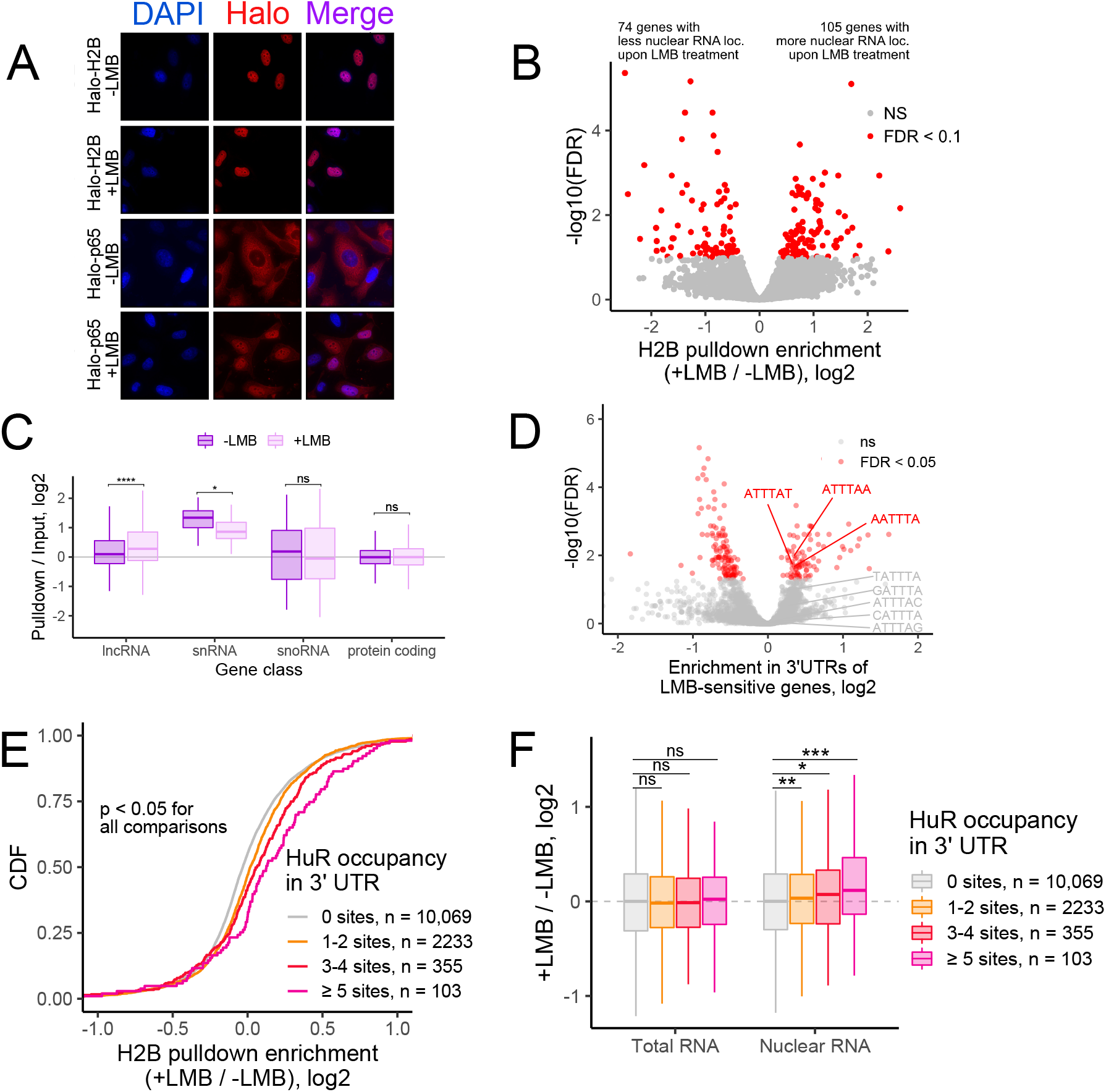
Halo-seq identifies transcriptome-wide changes in RNA localization following treatment with leptomycin B. (A) Visualization of the subcellular localization of Halo fusion proteins using a fluorescent Halo ligand. Note that while the p65 fusion is normally cytoplasmic, upon treatment with LMB, it becomes nuclear, indicating inhibition of CRM1 mediated export upon LMB treatment. (B) Volcano plot depicting changes in Halo-seq H2B pulldown enrichment (i.e. nuclear localization) following LMB treatment. (C) Changes in H2B pulldown enrichment following LMB treatment for selected gene classes. (D). Kmers of length 6 enriched in the 3′ UTRs of genes whose H2B pulldown enrichment was sensitive to LMB compared to the 3′ UTRs of genes whose H2B pulldown enrichment was not sensitive to LMB. (E) Genes were binned by the number of HuR binding sites (as defined by CLIP-seq) in their 3′ UTRs. Enrichments in H2B pulldown samples from LMB treated and untreated samples were then compared. (F) Abundance changes in total RNA (left) and nuclear RNA (right) following LMB treatment as a function of the number of HuR binding sites in the 3′ UTRs of transcripts. Total RNA and nuclear RNA samples correspond to the input and pulldown samples from H2B-targeted Halo-seq experiments. All significance tests were performed using a Wilcoxon rank-sum test. p value notation: * < 0.05, ** < 0.01, *** < 0.001, **** < 0.0001.

LMB treated and untreated samples displayed approximately the same amount of nuclear RNA as assayed by streptavidin-HRP RNA dotblot (**Figure S4A**). RNAseq analysis of the treated and untreated samples revealed that the RNA of 105 genes was significantly more enriched in the H2B pulldown sample in the LMB treated sample than the LMB untreated sample (FDR < 0.1) (**Figure 5B, S4B, Table S4**).

We then asked about the relative nuclear enrichment of different RNA classes (**Figure 5C**). As a class, lncRNAs were more nuclearly enriched following LMB treatment, suggesting that they may broadly depend on CRM1 for export. The nuclear enrichment of snRNAs was decreased following LMB treatment, in line with reports documenting that CRM1 is required for snRNP maturation and retention in nuclear Cajal bodies (Sleeman, 2007). The nuclear enrichment of snoRNAs and protein-coding mRNAs were unaffected.

We defined transcripts whose nuclear export was sensitive to LMB (FDR < 0.05) using the software package Xtail (Xiao et al., 2016). To identify transcript features associated with LMB sensitivity, we used a software package we created called FeatureReachR (https://github.com/TaliaferroLab/FeatureReachR). Using FeatureReachR, we found that the 3′ UTRs of transcripts whose localization was LMB-sensitive were enriched for several AU-rich kmers (**Figure 5D, S4C**).

Given the enrichment observed in LMB-sensitive transcripts for AU-rich kmers and motifs, we then directly searched these transcripts for AREs using a previously compiled database of their locations (Bakheet et al., 2018). We found that LMB-sensitive transcripts were enriched for AREs in their 3′ UTRs (**Figure S4D**).

AREs can be bound by several different RBPs, including HuR (also known as ELAVL1) (Peng et al., 1998). HuR has previously been identified as a protein whose export to the cytoplasm depends on CRM1 (Gallouzi et al., 2001). Using FeatureReachR’s ability to search for enrichment of the known binding motifs of many RBPs, we found that HuR RNA motifs were enriched in the 3′ UTRs of LMB-sensitive transcripts (**Figure S4E**). To extend this analysis to HuR binding sites identified in cells, we used HuR CLIP-seq data. The number of HuR binding sites in the 3′ UTRs of LMB-sensitive transcripts showed a dose-dependent relationship with the amount of increased nuclear enrichment of the transcript following LMB treatment (**Figure 5E**).

Since Halo-seq enrichments are calculated as a ratio of abundances in streptavidin-pulldown and input RNA, the increased enrichment of HuR-bound RNAs following LMB treatment could be due to either an increase in the abundance of HuR targets in the H2B pulldown or a decrease in their abundance in input samples. To distinguish between these possibilities, we compared LMB-induced abundance changes in the input and H2B pulldown samples separately. We found that the abundance of HuR target RNAs in total RNA (input) samples did not change upon LMB treatment. In contrast, the abundance of HuR target RNAs in the nuclear (pulldown) samples significantly increased upon LMB treatment (**Figure 5F**). These results suggest that hundreds of RNAs depend on HuR for efficient export from the nucleus.

## DISCUSSION

In this study, we have developed an RNA proximity method called Halo-seq that has the capability to isolate and quantify subcellular transcriptomes. We used Halo-seq to analyze nuclear and cytoplasmic transcriptomes and found that ARE-containing transcripts are relatively enriched in the nucleus and depleted from the cytoplasm. The capacity of AREs to negatively regulate RNA stability has been extensively established (Mukherjee et al., 2011; Shaw and Kamen, 1986), but their ability to induce differential nucleocytoplasmic localization at steady state has not been reported. One of the key RBPs that binds AREs to induce RNA degradation, Tristetraprolin, is predominantly localized to the cytoplasm (Johnson et al., 2002). One possible explanation, then, of the observed nuclear enrichment and cytoplasmic depletion of ARE-containing transcripts is that these RNAs are more vulnerable to degradation in the cytoplasm than in the nucleus (**Figure 6A**).

**Figure 6.**
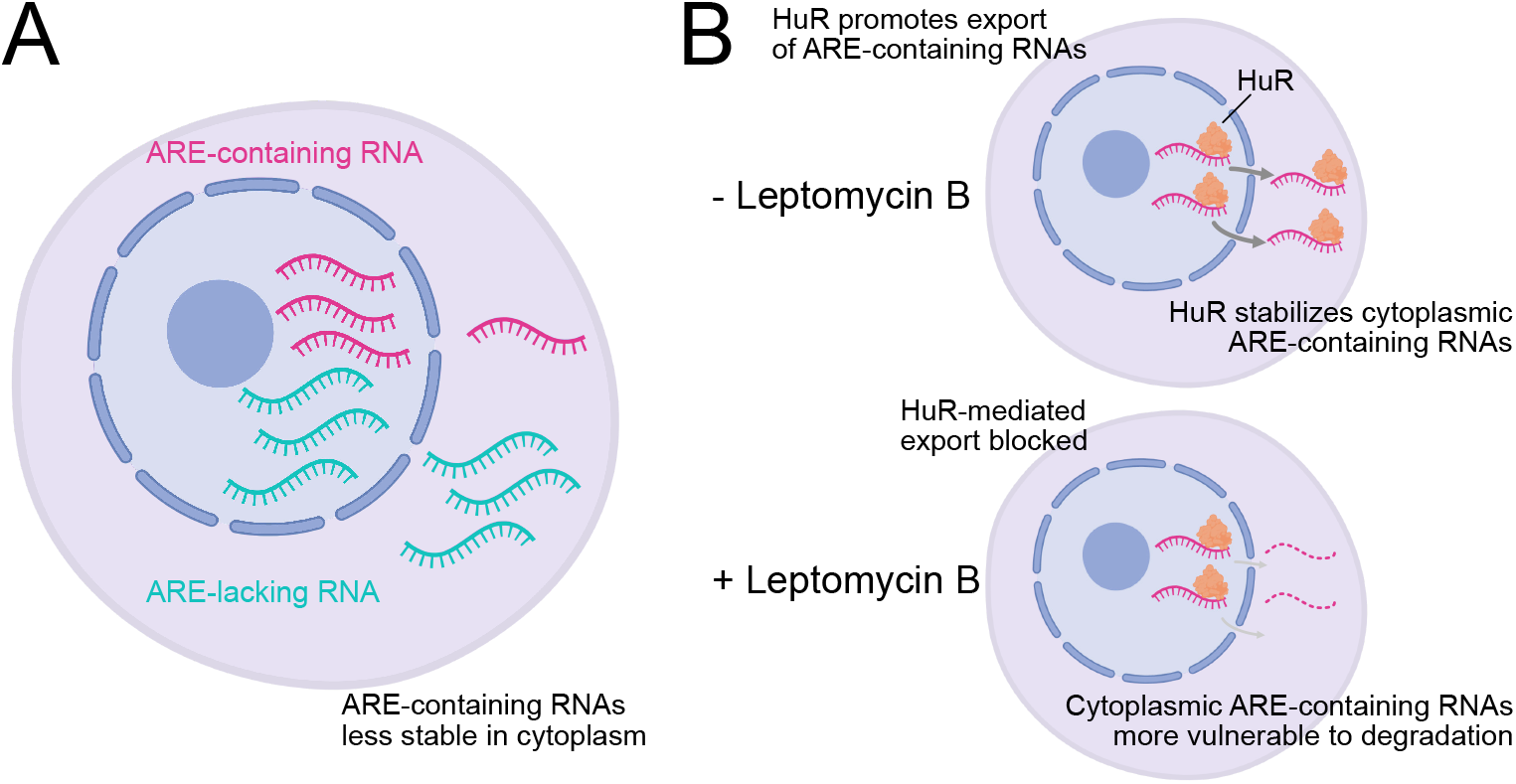
Model for observed differences in nuclear and cytoplasmic transcript abundances. (A) ARE-containing RNAs were relatively less abundant in the cytoplasm than the nucleus in our Halo-seq experiments. Cytoplasmic localization of many proteins that drive ARE-mediated RNA degradation, including TTP, could be a major contributing factor to this observation. (B) Upon LMB treatment, ARE-containing RNAs become even more relatively enriched in the nucleus. AREs are often bound by HuR, which both participates in RNA export and stabilizes cytoplasmic ARE-containing transcripts. HuR export from the nucleus is blocked by LMB, resulting in decreased export of ARE-containing transcripts and/or their increased vulnerability to degradation in the cytoplasm.

We then assayed dynamic RNA localization in response to treatment with the nuclear export inhibitor LMB. We found a dose-dependent relationship between the number of 3′ UTR HuR binding sites an RNA contained and the degree to which its abundance in the nucleus was increased following LMB treatment. HuR is known to both increase the stability of the RNAs that it binds (Mukherjee et al., 2011) and be important for the nuclear export of at least a handful of RNAs (Gallouzi et al., 2001). Although HuR shuttles between the nucleus and cytoplasm, its export from the nucleus is inhibited by LMB (Gallouzi et al., 2001). We therefore propose two non-exclusive models to explain the observed increase in nuclear abundance of HuR bound RNAs following LMB treatment (**Figure 6B**).

In one scenario, LMB-induced retention of HuR in the nucleus increases the vulnerability of its target RNAs to cytoplasmic degradation as its ability to protect its targets from instability has been lost. In an alternative scenario, RNAs that depend on HuR for nuclear export are now less efficiently exported, leading to their accumulation in the nucleus.

Nuclear RNA makes up a relative minority of total cellular RNA while cytoplasmic RNA makes up the majority. If the primary effect of LMB treatment on HuR-bound transcripts was a loss of stability, particularly in the cytoplasm, then we would expect to find that their levels in total RNA would be decreased. However, if the loss of HuR-mediated export was heavily involved, then we would expect to see HuR-bound RNAs become more abundant in the nucleus. Since, though, nuclear RNA makes up a small fraction of total RNA, their overall abundance may be unchanged. Importantly, we found that the overall levels of HuR-bound transcripts were unchanged following LMB treatment while their levels in the nucleus significantly increased (**Figure 5H**). These results are consistent with the idea that HuR regulates the nuclear export of hundreds of transcripts.

We believe these results are important in the context of the existing literature describing HuR activity. Although HuR has been previously found to regulate the nuclear export of a limited number of RNAs (Gallouzi et al., 2001), the vast majority of the work on HuR has focused on its capacity to regulate RNA stability. These results therefore simultaneously greatly expand our knowledge of the transcripts that use HuR for nuclear export and provide strong confidence that Halo-seq can accurately quantify dynamic subcellular localization and be used to derive mechanistic insights regarding its regulation.

Using known markers of nuclear and nucleolar RNA, we found that Halo-seq can efficiently distinguish the RNA contents of these two compartments. Given their close spatial proximity, these results favorably report on the ability of Halo-seq to quantify RNA localization with high spatial precision.

The ability of Halo-seq to recapitulate known RNA localization patterns as well as its ability to connect these patterns to activities of specific RBPs gave us confidence that the method performed well. However, a number of approaches for quantifying subcellular transcriptomes had been previously reported (Benoit Bouvrette et al., 2018; Fazal et al., 2019; Kaewsapsak et al., 2017; Padrón et al., 2019; Wang et al., 2019). We therefore compared the efficiency and specificity of RNA labeling produced by Halo-seq to these methods.

CeFra-seq uses biochemical fractionation to define subcellular fractions (Benoit Bouvrette et al., 2018). Although we observed some agreement between the nuclear and cytoplasmic RNA contents defined by CeFra-seq, we found that Halo-seq performed better in terms of correctly placing RNAs of known localization. A key advantage of proximity labeling schemes, including Halo-seq, over biochemical fractionations is that the important labeling steps are performed while cells are alive and intact. Biochemical fractionations are inherently susceptible to false positives in which RNAs that are spatially distinct within a cell yet possess similar biochemical properties copurify. They are also susceptible to false negatives in which the association of spatially coincident RNAs does not survive the biochemical purification. These factors may explain the superior performance of Halo-seq.

Two RNA proximity labeling techniques, APEX-seq and CAP-seq, were also recently reported (Fazal et al., 2019; Padrón et al., 2019; Wang et al., 2019). We observed a general agreement in the quantification of localized RNAs with Halo-seq and APEX-seq, although we were unable to quantify the localization of small RNAs in the APEX-seq data due to their use of oligo-dT enrichment instead of rRNA depletion. We were unable to directly compare RNA localization patterns observed by Halo-seq to those observed by CAP-seq due to the fact that the raw sequencing data from the report describing CAP-seq has not to our knowledge been made publicly available.

We compared RNA labeling efficiencies of all three proximity labeling methods (**Figure 4B**,**C**). We found that Halo-seq was significantly more efficient in generating labeled RNA than both APEX-seq and CAP-seq. These observations may be attributed to differences in the efficiency of singlet oxygen generation (as compared to miniSOG) or the reactive species that is being generated in comparison to the singlet oxygen from DBF excitation.. The increased efficiency of Halo-seq may make it better suited to the study of precisely defined subcellular RNA populations that make up a relatively small proportion of the total cellular RNA content. Halo-seq may therefore allow interrogation of the RNA content of small, defined locations that until now were intractable for such experiments. In sum, we view Halo-seq as a flexible, quantitative tool that is well-suited to advance the study of subcellular RNA localization.

## ACKNOWLEDGEMENTS

We thank members of the Taliaferro lab for helpful discussions regarding experiments and analyses. We also thank Neel Mukherjee for helpful discussion and tips regarding analysis of Halo-seq data.

This work was funded by the National Institutes of Health (R35-GM133885 to J.M.T.; DP2 GM119164 to R.C.S.) and the RNA Bioscience Initiative at the University of Colorado Anschutz Medical Campus (J.M.T. and R.G.). It was further supported by a Predoctoral Training Grant in Molecular Biology (NIH-T32-GM008730) (H.Y.G.L. and R.G.).

## METHODS

### Creation of transgenic cell lines expressing Halo fusion proteins

HeLa cell lines were created by integrating a plasmid containing the Halo fusion using cre/lox recombination (Khandelia et al., 2011). HeLa cells containing a single loxP cassette were plated in a 6-well plate and co-transfected with 2000 ng of a plasmid containing the Halo fusion and a puromycin selectable marker and 100 ng of a plasmid, pBT140, that expressed Cre recombinase. pBT140 was a gift from Liqun Luo (Addgene plasmid # 27493; http://n2t.net/addgene:27493; RRID:Addgene_27493) (Miyamichi et al., 2010).

The transfection was carried out using Lipofectamine 2000 (Thermo Scientific) according to the manufacturer’s instructions. Forty-eight hours after transfection, the cells were incubated with 5 µg / mL puromycin in order to select for integrants. Approximately 2 weeks after transfection, a stable population of integrants had been selected.

The plasmid containing the Halo fusion also contained a reverse tetracycline-controlled transactivator (rtTA), allowing the expression of Halo fusions to be controlled in a doxycycline-dependent manner. To induce expression of the Halo fusion, selected cell lines were incubated with 1 µg / mL doxycycline for 48 hours prior to performing experiments.

The H2B fusion was expressed as a C-terminal Halo fusion (i.e. H2B-Halo) while the p65 and fibrillarin fusions were expressed as N-terminal Halo fusions (i.e. Halo-p65 and Halo-fibrillarin).

### Validation of subcellular localization of Halo fusion proteins

To verify that the transgenic Halo fusion proteins were targeted to the correct subcellular location, the location of the fusions were visualized using a fluorescent Halo ligand. Cells were seeded on poly-D-lysine-coated coverslips and incubated with 1 µg / mL doxycycline for 48 hours. The media was then removed and the cells were washed one time with PBS. Cells were then fixed by incubating them in 2% formaldehyde for 30 minutes at room temperature. The cells were then washed with PBS again, and fluorescent Halo ligand (Janiela Fluor 646 (Promega), 25 nM in PBS) was added. The cells were incubated in the Halo ligand solution for 30 minutes at room temperature then washed three times with PBS for 5 minutes each. DAPI was then added to a concentration of 100 ng / mL for 10 minutes, and the cells were washed again with PBS. The coverslips were then mounted and imaged, usually at 60X magnification, using a Deltavision Elite widefield fluorescence microscope (GE).

### In-cell alkynylation of Halo-proximal RNAs

Cells were grown in 10 cm or 15 cm dishes, and the expression of Halo fusion proteins was induced with 1 µg / mL doxycycline for 48 hours. Cells were then washed with PBS and incubated with 1 µM DBF halo ligand (in HBSS) at 37°C for 15 minutes. For negative control samples, the DBF halo ligand was omitted.

Cells were then washed with complete media and incubated at 37°C twice for 10 minutes each. Propargylamine (Sigma, 1 mM in HBSS) was then added, and the cells were incubated for 5 minutes at 37°C. Following the incubation, cells were then irradiated with green light for 5 minutes from a 100 W LED flood light (USTELLAR). Dishes were sandwiched between two flood lights in a dark enclosed space so that they were illuminated from above and below. The light above the dish was suspended approximately 10 cm from the dish. Total RNA was then isolated from the cells using Trizol (Ambion) following the manufacturer’s instructions with the addition of a homogenization step in which the cells in Trizol solution are repeatedly forced through a 20 gauge needle 10 times.

The RNA was then DNase treated using DNase I (Thermo Scientific) for 30 minutes at 37°C. RNA was then again recovered using Trizol, and the final pellet was resuspended in water to a concentration higher than ∼300ng/ul. This ensures the ability to have the correct concentration in Click reactions downstream. Typical total RNA yields were 25-100 µg if starting with a 10 cm dish and 300-700 µg if starting with a 15 cm dish.

### In vitro biotinylation of alkynylated RNAs using Click chemistry

Typically, approximately 50-100 µg of total RNA was used in the Click reaction. The Click reaction contained 10 mM Tris pH 7.5, 2 mM biotin azide (Click Chemistry Tools), 10 mM sodium ascorbate made fresh (Sigma), 2 mM THPTA (Click Chemistry Tools), and 100 µM copper sulfate. The reaction was incubated for 30 min in the dark at 25°C. The reaction was then cleaned up using a Quick RNA Mini kit (Zymo Research) for small scale biotinylation tests (<10ug RNA), or through standard ethanol precipitation with sodium acetate for larger amounts of RNA (>10ug). RNA was then eluted and resuspended to 1 µg / µL in 50 mM NaCl.

### Streptavidin purification of biotinylated RNA

Typically, between 50 and 100 µg of RNA at 1 µg / µL from the Click reaction was then carried forward for streptavidin pulldowns. 25-50 µL of streptavidin-coated magnetic beads (Pierce PI88816) were used depending on the known amount of labeled RNA present in the reaction. Specifically, we used 1ul beads per 1ug total RNA for Halo-p65 and Halo-fibrillarin, and 1ul beads: per 2ug RNA for H2B. The beads were washed 3 times in B&W buffer (5 mM Tris pH 7.5, 0.5 mM EDTA, 1M NaCl, 0.1 Tween 20), 2 times in solution A (0.1 M NaOH, 50 mM NaCl), and 1 time in solution B (100 mM NaCl). The beads were resuspended in an appropriate amount of NaCl to allow for 50mM final concentration after the addition of the desired amount of RNA to equal 1 ug / ul. The RNA from the Click reaction was then added to the beads, and the beads were rotated for 2 hrs at 4°C. The beads were then washed 3 times for 5 minutes each in B&W buffer with rotation at room temperature.

RNA was then recovered from the beads using Trizol. First, the beads were resuspended in 50 µL PBS. 150 µL of Trizol was then added, and this mixture was incubated for 5-10 minutes at 37°C. The eluted RNA (without the magnetic beads) was then recovered from this mixture using a DirectZol kit (Zymo Research) following the manufacturer’s instructions and eluted in 10 µL of water. Depending on the location of the Halo fusion protein, typically between 0.5% and 5% of the input RNA was recovered by streptavidin pulldown.

### RNA dot blot

RNA biotinylation, both before and after the streptavidin pulldown, was assessed using an RNA dotblot. A 5 cm x 5 cm piece of Hybond-N+ membrane (GE) was wet with 2X SSC for 1 minute. This was then allowed to dry for 15 minutes. RNA samples, typically around 5 µg, were then spotted on the membrane and allowed to dry for 30 min. The dried blots were then crosslinked twice to the membrane using 120,000 µJ / cm2 on a Stratalinker UV crosslinker (Stratagene).

To stain for total RNA, the blot was then incubated in 1% methylene blue for 10 minutes and destained using deionized water. The membrane was then blocked using 5% BSA for 30 minutes and washed 3 times in PBST (PBS + 0.01% Tween). Biotinylated RNA was then detected using streptavidin-HRP (Abcam ab7403) at a dilution of 1 to 20,000 in 3% BSA by addition to the membrane with rocking overnight at 4°C. The membrane was then washed 3 times for 10 min each in PBST at room temperature. Streptavidin-HRP was then detected using standard HRP chemiluminescent reagents (Advansta) and visualized using chemiluminescent imaging on a Sapphire molecular imager (Azure Biosystems).

### Library preparation and high-throughput sequencing

rRNA-depleted RNAseq libraries were prepared using an RNA HyperPrep Kit (KAPA / Roche). 100 ng of RNA were put into the beginning of the library prep protocol, and 14 PCR cycles were used to amplify the library at the end.

Libraries were sequenced using paired end sequencing (2 × 150 bp) on a NovaSeq high-throughput sequencer (Illumina) at the University of Colorado Genomics Core Resource. Typically, between 20 and 40 million read pairs were sequenced for each sample.

### Analysis of RNAseq data to identify genes enriched in streptavidin pulldown

Transcript abundances in RNAseq data were quantified using Salmon (Patro et al., 2017) and a human genome annotation retrieved from GENCODE (www.gencodegenes.org, GENCODE 28). Gene abundances were then calculated from these transcript abundances using tximport (Soneson et al., 2015), and genes whose abundance in input and streptavidin-pulldown samples were identified using DESeq2 (Love et al., 2014).

### Analysis of unspliced transcripts in RNAseq data

In order to quantify the relative abundances of spliced and unspliced (all introns remaining) versions of transcripts, a custom fasta file was supplied to Salmon which contained two versions of every transcript, one with all introns remaining and one with all introns removed. The custom fasta file was generated using this script: https://github.com/rnabioco/rnaroids/blob/master/src/add_primary_transcripts.py. Salmon then assigned reads competitively to these transcripts. For each gene, the ratio of abundances of unspliced and spliced transcripts was then calculated.

### Definition of AU-rich elements within transcripts

AU-rich element locations were downloaded from the AU-rich element database (Bakheet et al., 2018). AU-rich elements within 3′ UTRs were used for analysis.

### In situ Click reaction using Cy5-azide

To visualize the subcellular location of DBF-proximal alkynylated molecules in situ, cell growth and Halo fusion induction were performed as described above with the exception that cells were grown on poly-D-lysine-coated coverslips. The in-cell alkynylation of DBF-proximal molecules was also performed as described above. Then, instead of lysing cells with Trizol to recover RNA, cells were washed with PBS 3 times and fixed by incubating them in fixation buffer (3.7% formaldehyde, 0.1% Triton in PBS) for 30 minutes at room temperature. Cells were then washed twice for 5 minutes each in PBS.

The Click reaction was then performed by incubating cells with 100 µL Click buffer (100 µM copper sulfate, 2 mM THPTA, 10 mM fresh sodium ascorbate, 10 µM Cy5 picolyl azide (Click Chemistry Tools 1171-1)) for 1 hour at 37°C in the dark. As two separate controls, samples in which the DBF had been left out during the alkynylation reaction and the Cy5 had been left out of the Click buffer were used. Following the Click reaction, the coverslips were washed three times for 5 minutes each in wash buffer (0.1% Triton, 1 mg / mL BSA in PBS). Coverslips were then incubated in DAPI buffer (100 ng / mL DAPI in wash buffer) for 30 minutes at 37°C and then washed twice with wash buffer for 5 minutes each. The coverslips were then mounted and imaged using a Deltavision Elite widefield fluorescence microscope (GE).

### Treatment with Leptomycin B

Cells were treated with 40 ng / mL leptomycin B (LMB) for 15 hours prior to the beginning of the Halo-seq protocol. To verify the activity of leptomycin B and our conditions, the localization of the Halo-p65 fusion was monitored. The cytoplasmic localization of p65 is sensitive to LMB (Wolff et al., 1997) (**Figure 5A**). Accordingly, the normally cytoplasmically localized Halo-p65 fusion became localized to the nucleus following LMB treatment (**Figure 5A**).

### Identification of RNAs whose subcellular localization was sensitive to LMB

Halo-seq using a histone H2B Halo fusion was performed in the presence and absence of LMB, generating four conditions: input samples with and without LMB and pulldown samples with and without LMB. To identify genes whose pulldown to input ratio changed between the LMB treated and untreated samples, the software package Xtail was used (Xiao et al., 2016). Xtail is designed for the analysis of ribosome profiling data and identification of genes whose ribosome occupancy changes across conditions. Ribosome profiling data is the ratio of two gene expression values, one drawn from ribosome footprints and the other drawn from bulk RNA. Xtail identifies genes whose ratio changes between two conditions. The analysis of Halo-seq data is structurally similar as it is also a ratio of gene expression values, one from an input sample and another from a pulldown sample. We therefore used Xtail to identify genes whose ratio (i.e. nuclear enrichment) changed between LMB treated and untreated samples.

### Comparison of RNA labeling with miniSOG2

Cells stably expressing HA-miniSOG2 were treated alongside cells stably expressing Halo-HA. The expression of miniSOG2 and Halo-HA was induced with 1 µg / mL doxycycline for 48 hours. Cells were then washed with PBS. Halo-expressing cells were then incubated with Halo-DBF ligand at 37°C for 5 minutes, and both cell lines were then incubated with propargylamine (Sigma, 1mM in HBSS) at 37°C for 5 minutes. Following the incubation, cells were irradiated with blue light (for miniSOG2) or green light (for Halo-DBF) for a total of 10 minutes with a 100 W LED flood light (USTELLAR). Dishes were sandwiched between two flood lights so that they were illuminated from above and below. The light above the dish was suspended approximately 10 cm from the dish. Total RNA was then isolated from the cells using Trizol (Ambion) following the manufacturer’s protocol after homogenization with a 20 gauge needle 10 times. RNA was biotinylated following the protocols described above.

### Comparison of RNA labeling with APEX2

A cell line was created expressing a Halo-APEX2 fusion protein to ensure equal expression of HaloTag domain and APEX2 protein. RNA labeling with APEX2 was done alongside Halo-DBF-mediated labeling (described above). Expression of the Halo-APEX2 fusion protein was induced with 1 µg / mL of doxycycline for 48 hours. To label RNA with the APEX2 protein, cells were washed with PBS and then incubated with HBSS containing 0.5mM biotin phenol (APExBIO A8011) for 30 minutes at 37°C. Cells were then washed in PBS and either treated with hydrogen peroxide at 1mM in HBSS or plain HBSS (control) for 7 minutes at 37°C (labeling of compared RNA using the Halo-DBF method was also subject to 7 minutes of exposure to green light). The reaction was quenched in quenching buffer containing 5mM Trolox, 10mM sodium ascorbate, and 10mM sodium azide in PBS. To fully quench the reaction, cells were washed three times in quenching buffer, with 3 minutes between each wash. Total RNA was isolated from the cells using Trizol (Ambion) after homogenization with a 20 gauge needle. After DNAse treatment, RNA was then directly blotted onto an RNA dotblot.

### Comparison with APEX-seq data

APEX-seq RNAseq data was downloaded from the Gene Expression Omnibus (GSE116008) and processed to calculate transcript and gene abundances as above. Since the libraries were produced using poly-A enrichment of RNA, only protein-coding and lncRNA genes were used for comparisons to Halo-seq data.

### Comparison with CeFra-seq data

CeFra-seq RNAseq data was downloaded from the ENCODE portal (www.encodeproject.org). Transcript and gene abundances from rRNA-depleted libraries were calculated using Salmon, tximport, and DESeq2 as outlined above. While Halo-seq enrichments were calculated as a gene’s abundance in the streptavidin pulldown divided by its abundances in the input to the pulldown, CeFra-seq enrichments were calculated as a gene’s abundance in the biochemically defined fraction (e.g. cytosol or nucleus) divided by its abundance in the total RNA samples.

## DATA AVAILABILITY

All high-throughput RNA sequencing data as well as transcript quantifications have been deposited at the Gene Expression Omnibus under accession number GSE172281.

## SUPPLEMENTARY FIGURES

**Figure S1.**
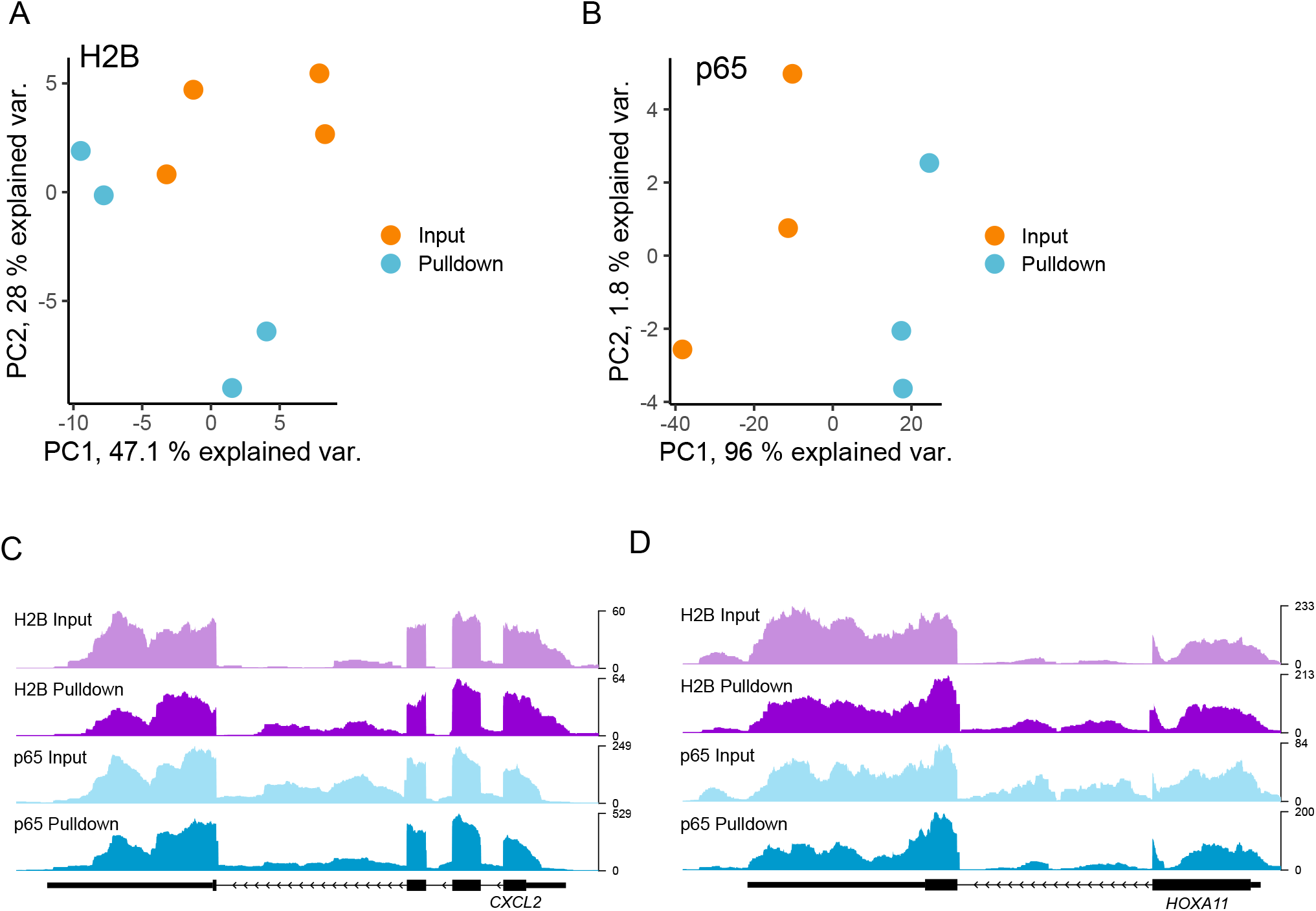
(A) Principal component analysis of gene expression values from the Halo-seq histone H2B experiment. (B) Principal component analysis of gene expression values from the Halo-seq p65 experiment. (C) Read coverage over *CXCL2* in Halo-seq histone H2B and p65 experiments. Note the higher intronic coverage in the H2B pulldown sample relative to the H2B input sample, whereas the p65 pulldown sample has less intronic coverage relative to the p65 input sample. (D) As in C, but for the gene *HOXA11*.

**Figure S2.**
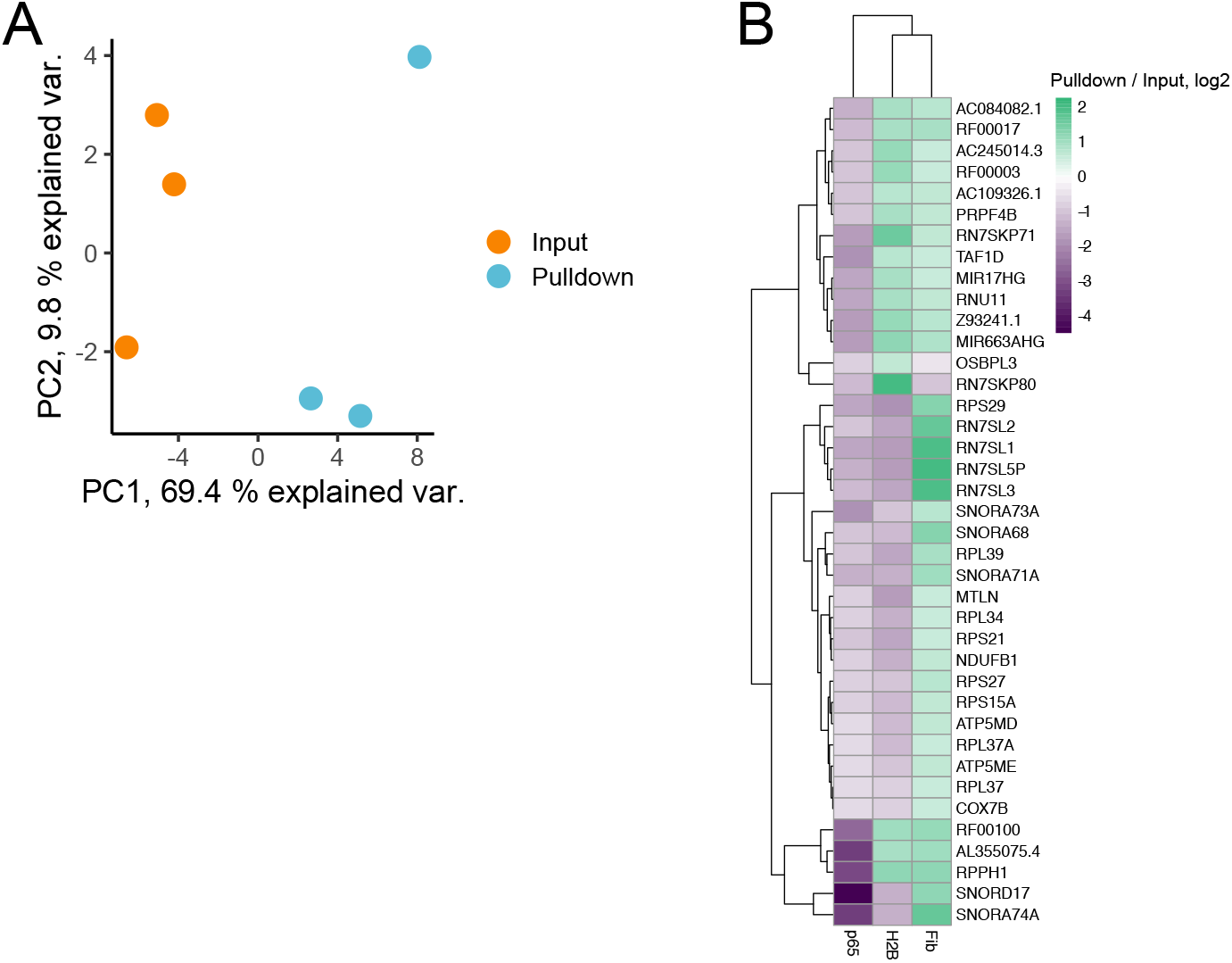
(A) Principal component analysis of gene expression values from the Halo-seq fibrillarin experiment. (B) Selected genes whose RNAs were specifically enriched in the Halo-seq fibrillarin experiment and not the histone H2B or p65 experiments.

**Figure S3.**
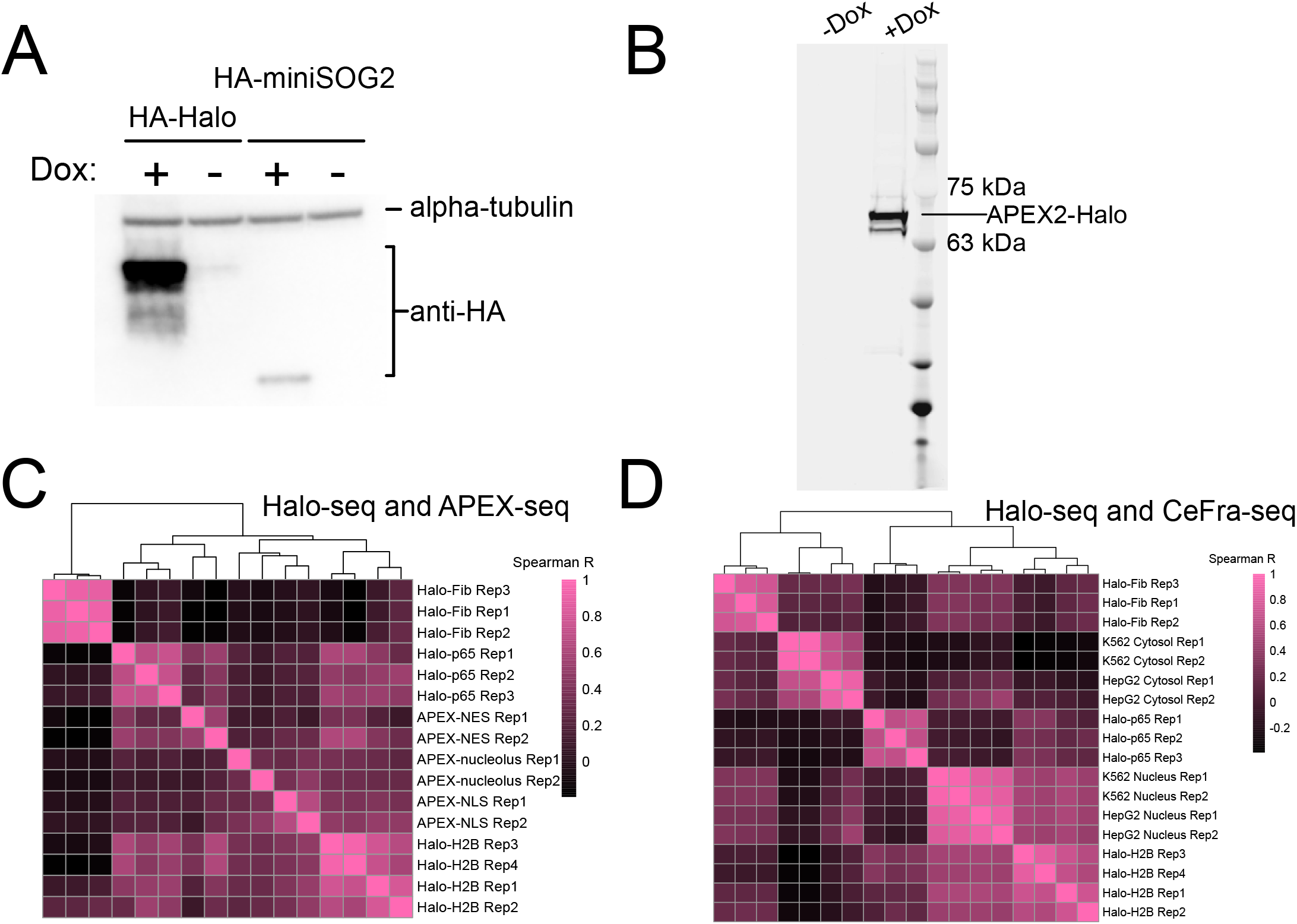
(A) Expression of Halo and minisog2 proteins in HeLa cells. Both proteins were HA tagged, and their expression was inducible with doxycycline. (B) Expression of APEX2-Halo fusion in HeLa cells. Halo fusion proteins were visualized through the addition of fluorescent Halo ligand to lysate prior to electrophoresis. (C) Spearman correlation of gene-wise enrichment values (streptavidin pulldown / input) for Halo-seq and APEX-seq samples. (D) Spearman correlation of gene-wise enrichment values (streptavidin pulldown / input for Halo-seq; biochemical fraction / total for CeFra-seq) for Halo-seq and CeFra-seq samples.

**Figure S4.**
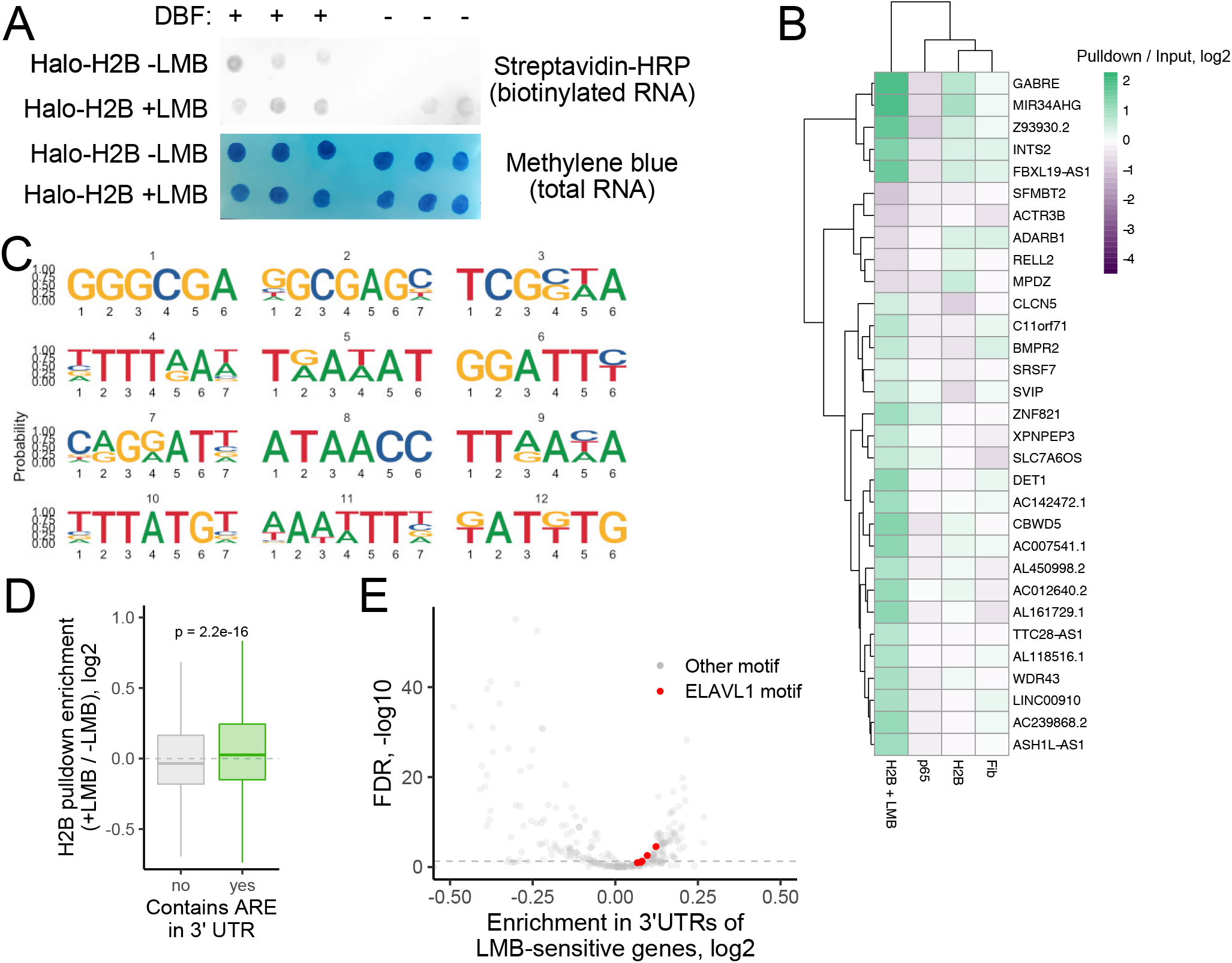
(A) RNA biotinylation, as assayed by RNA dot blot on Halo-seq histone H2B samples with and without LMB treatment. (B) Selected genes whose RNAs were specifically enriched or depleted in the Halo-seq histone H2B experiment following LMB treatment. (C) RNA sequence motifs enriched in the 3′ UTRs of genes that become more enriched in the nucleus following LMB treatment. (D) Changes in H2B pulldown enrichment upon LMB treatment for genes that either do or do not contain AU-rich elements in their 3′ UTRs. (E) RBP binding motif enrichment in the 3′ UTRs of genes that become more enriched in the nucleus following LMB treatment. RBP binding motifs were defined using data from RNA bind-n-seq (Dominguez et al., 2018), and their enrichments were calculated using FeatureReachR.

**Table S1**. RNAseq enrichments (pulldown / input) for the Halo-p65 Haloseq experiment, as identified by DESeq2. Gene biotype designations are drawn from biomaRt.

**Table S2**. RNAseq enrichments (pulldown / input) for the H2B-Halo Haloseq experiment, as identified by DESeq2. Gene biotype designations are drawn from biomaRt.

**Table S3**. RNAseq enrichments (pulldown / input) for the Fibrillarin-Halo Haloseq experiment, as identified by DESeq2. Gene biotype designations are drawn from biomaRt.

**Table S4**. Changes in H2B-Halo pulldown enrichment following LMB treatment. Log2FC columns represent the genes enrichment in the H2B-Halo pulldown sample compared to input samples. Enrichment change represents the change in this enrichment between LMB treated and untreated samples (treated - untreated). The p value column has been adjusted for multiple hypothesis testing and is the result of tests asking whether the treated and untreated log2FC values were different. This table was modified from the output of Xtail software (Xiao et al., 2016).

